# Polymorphisms in immunoglobulin heavy chain variable genes and their upstream regions

**DOI:** 10.1101/2020.01.27.921197

**Authors:** Ivana Mikocziova, Moriah Gidoni, Ida Lindeman, Ayelet Peres, Omri Snir, Gur Yaari, Ludvig M. Sollid

## Abstract

Germline variations in immunoglobulin genes influence the repertoire of B cell receptors and antibodies, and such polymorphisms may impact disease susceptibility. However, the knowledge of the genomic variation of the immunoglobulin loci is scarce. Here, we report 25 novel germline *IGHV* alleles as inferred from rearranged naïve B cell cDNA repertoires of 98 individuals. Thirteen novel alleles were selected for validation, out of which ten were successfully confirmed by targeted amplification and Sanger sequencing of non-B cell DNA. Moreover, we detected a high degree of variability upstream of the V-region in the 5’UTR, leader 1, and leader 2 sequences, and found that identical V-region alleles can differ in upstream sequences. Thus, we have identified a large genetic variation not only in the V-region but also in the upstream sequences of *IGHV* genes. Our findings challenge current approaches used for annotating immunoglobulin repertoire sequencing data.

---

Immunoglobulins are an important part of the adaptive immune system. They exert their function either as the antigen receptor of B cells that is essential for the antigen presentation capacity of these cells, or as secreted antibodies that survey extracellular fluids of the body. Immunoglobulins can bind a plethora of antigen epitopes via their paratopes, which are composed of combinations of heavy and light chain’s variable regions. A huge diversity of paratopes is established by recombination of variable (V), diversity (D) (not in light chains) and joining (J) genes, and the pairing of heavy and light chains^1^. There is a large number of V, D, and J genes present on the heavy chain locus (chromosome 14, 14q32.33)^2^ as well as the two light chain loci kappa (chromosome 2, 2p11.2) and lambda (chromosome 22, 22q11.2)^3^.

These loci remain incompletely characterized due to the fact that they contain many repetitive sequence segments with many duplicated genes^4^, which makes it difficult to correctly assemble short reads from whole genome sequencing. Single nucleotide polymorphisms as well as copy number variations are in linkage disequilibrium and make up distinct haplotypes^4^. To this date, a limited number of genomically sequenced ^5–7^ and inferred ^8, 9^ haplotypes of the heavy chain and the two light chain loci have been described. Different databases exist for genomic immune receptor DNA sequences (IMGT/GENE-DB^10^), putative novel variants from inferred data (IgPdb^11^) or entire immune receptor repertoires (OGRDB^12^).

The usage of immunoglobulin heavy chain variable (*IGHV)* genes and their mutational status are most frequently studied in relation to cancer^13, 14^, responses to vaccines^15, 16^, or in autoimmune diseases^17–19^. Most *IGHV* genes have several allelic variants and more alleles are being discovered as a result of adaptive immune receptor repertoire-sequencing (AIRR-seq)^20, 21^. Software tools such as TIgGER^22, 23^, IgDiscover^24^ and partis^25^ allow to infer germline alleles from such repertoire data. Based on these inferred alleles, the data can then be input to other tools that infer haplotypes and repertoire deletions^26^. Incorrect annotation could possibly lead to inferring wrong deletions and biased assessments. Therefore, having a full overview of germline variants is essential for studying the adaptive immune response with high accuracy.

Some allelic variants have been associated with increased disease susceptibility^27, 28^, yet the impact of immunoglobulin gene variation on disease risks is still unknown^29^. These regions have not been sufficiently covered in the numerous genome wide association studies performed to date. More comprehensive maps of polymorphisms are required for proper analysis.

Here, we have used previously generated AIRR-seq data^30^ from naïve B-cells of 98 Norwegian individuals to identify novel *IGHV* alleles, a selection of which we then validated from genomic DNA (gDNA) of non-B cells, i.e. T cells and monocytes. We also analyzed the sequences upstream of the V-region, and constructed consensus sequences for the upstream variants present in the cohort. These results expand our knowledge of this important locus and deepen our understanding of allelic diversity within the Caucasian population. In addition, the result of this study can be used to improve the accuracy of currently used bioinformatics tools for the analysis of immunoglobulin repertoire sequencing data.

## RESULTS

In this study, we used an AIRR-seq dataset from a cohort of 98 individuals^30^ aiming to characterize novel *IGHV* alleles that might be present in the data, as well as analyze the sequences upstream of the V-region and create a table of previously unexplored upstream variants (Fig.1). To validate our inferences from the AIRR-seq data analysis, genomic DNA of the same individuals was isolated from non-B cells, i.e. T cells and monocytes. The reason for using non-B cells for validation was to avoid capturing sequences with somatic hypermutation that occurs in primed B cells.

**Figure 1.**
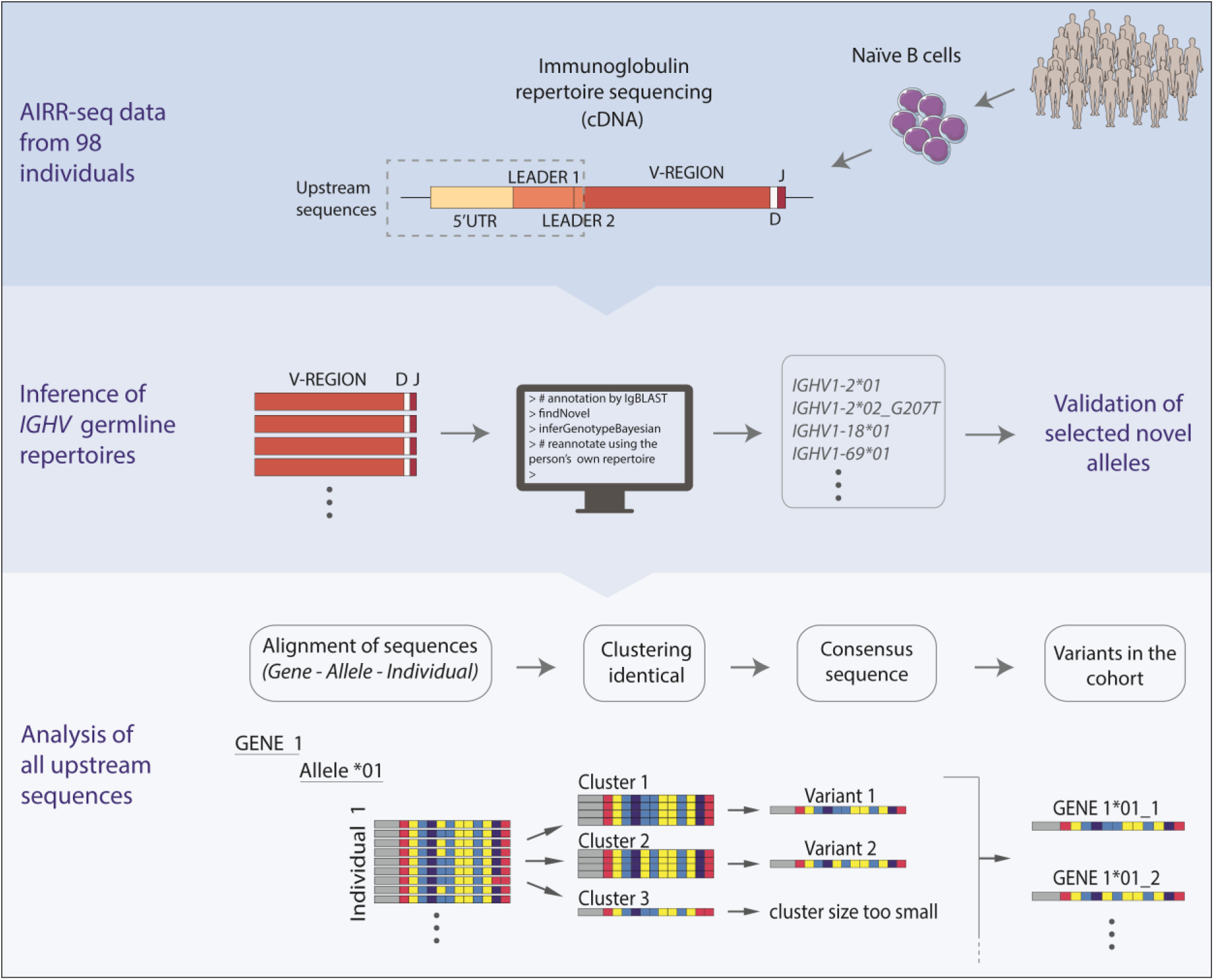
Schematic representation of the data analysis. In this study, we used material from a Norwegian cohort of 98 individuals^30^. Following the initial preprocessing of the data, we inferred the germline V-gene repertoires of all individuals in the cohort and identified novel alleles using the software suites TIgGER and IgDiscover. The availability of genomic DNA of the same individuals allowed us to verify some of our findings from the analysis of the AIRR-seq data. Since the validation attempts revealed polymorphisms outside of the V-region, we decided to analyze the upstream sequences, i.e. 5’UTR, leader 1 and leader 2. We used a custom approach for this analysis based on clustering identical variants. More details about the protocols and analysis can be found in the methods section.

We used two germline inference tools, TIgGER^22, 23^ and IgDiscover^24^, to characterize novel alleles and to create a personalized germline reference of *IGHV* alleles for each individual (aka genotype). The purpose of using two different software tools was to increase our confidence in the inference of novel alleles. This study does not aim to serve as a comparison of the available software tools.

To increase the overlap between the different software results and to allow the discovery of novel alleles in genes with low expression, we adjusted selected TIgGER parameters, while keeping the IgDiscover parameters as default. Suspected false positive signals were filtered out from the novel allele candidates using mismatch frequency as described in Methods. The mismatch frequencies are depicted in Supplementary Fig.1. Novel allele candidates that were determined to be false positives contained mutations A152G, T154G and A85C (Supplementary Fig.1).

### Analysis of the V-region reveals 25 novel *IGHV* alleles

We first analyzed the usage of all genes and the different alleles carried by individuals in the cohort. The relative usage of certain genes appeared to be strongly affected by the alleles present in the inferred genotype. This was true for *IGHV1-2, IGHV1-46, IGHV3-11, IGHV3-43, IGHV3-48, IGHV3-53, IGHV4-61*, and *IGHV5-51* (Supplementary Fig.2). Overview of the usage of all genes across all individuals can be found in Supplementary Fig.2-3.

We inferred altogether 25 novel alleles (Fig.2), and we named them using the closest reference allele. The majority of the novel alleles (22) were identified both with TIgGER and IgDiscover. In addition to these, two novel alleles were found exclusively by IgDiscover, and one novel allele was found exclusively by TIgGER.

**Figure 2.**
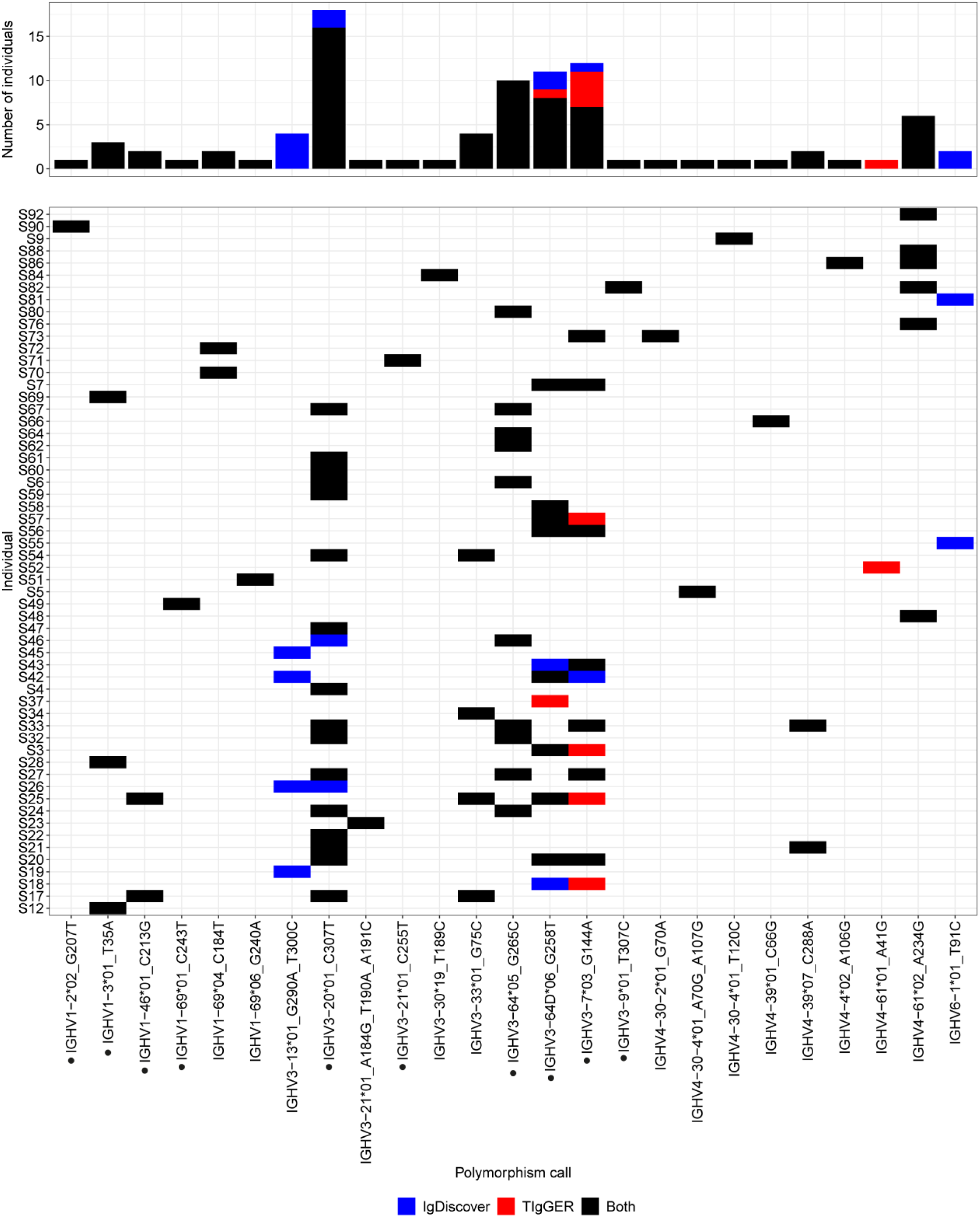
Novel *IGHV* alleles. The software suites TIgGER and IgDiscover were used to infer a personal *IGHV* genotype for each individual and to infer previously undiscovered alleles. All novel alleles that are part of a genotype inferred by at least one of the methods appear on the x-axis. Alleles that were validated by Sanger sequencing are marked with a dot. Individuals with at least one novel allele lie on the y-axis and are labelled by their subject name. For each allele, the color of a tile (or a bar) represents the method of detection and genotype inference. The height of each bar on top represents the number of individuals for whom a certain allele was inferred and is part of a genotype.

Thirteen novel alleles were selected for validation by targeted amplification and subsequent Sanger sequencing of gDNA (Supplementary Fig.4) of non-B cells, i.e.T cells and monocytes isolated by fluorescence-activated cell sorting^30^. The validation primers are specified in the Supplementary Table 1. Out of those thirteen alleles, ten were successfully validated. These include *IGHV1-2*02_G207T, IGHV1-3*01_T35A, IGHV1-46*01_C213G, IGHV1-69*01_C243T, IGHV3-7*03_G144A, IGHV3-9*01_T307C, IGHV3-20*01_C307T, IGHV3-21*01_C255T, IGHV3-64*05_G265C*, and *IGHV3-64D*06_G258T*. Surprisingly, *IGHV3-64*05_G265C* was found to originate from *IGHV3-64D* (Fig.6c). Two of the novel alleles, namely *IGHV1-46*01_C213G* and *IGHV3-20*01_C307T*, have been recently added to the IMGT database as *IGHV1-46*04* and *IGHV3-20*04* respectively.

Validation of the novel alleles revealed additional polymorphisms outside of the V-region. The allele *IGHV3-64*06_G258T* has a polymorphism in leader 1 (position −21) in addition to the V-region polymorphism. Genomic validation of *IGHV3-7*03_G144A* revealed a further polymorphism in the intron. During validation of this allele, we also managed to amplify the genomic sequence of *IGHV3-7*02*, which carried the previously reported polymorphism A318G^31^. This polymorphism was not inferred from the AIRR-seq data in our study, since the default parameters of the inference tools are set to detect polymorphisms up to position 312.

Attempts to validate *IGHV4-39*07_C288A*, *IGHV4-61*02_A234G*, and *IGHV6-1*01_T91C* were unsuccessful. The gene-specific primers that were used for validation were designed based on the current reference genome. However, the efficiency of the *IGHV4* primers was inferior, and Sanger sequencing only revealed allele *\*01* of each gene, even in clearly heterozygous individuals.

### Analysis of upstream sequences yields a more complete and accurate germline reference dataset

As some of the validated novel alleles had additional polymorphisms in the intron or the leader sequence, we extended our analysis of the AIRR-seq data beyond the V-region. Although introns are not present in the AIRR-seq data, the sequences of the 5’ untranslated region (5’UTR), leader 1, and leader 2 lie upstream of the V-region and are present in the data thanks to the library preparation method (Fig.1). We will refer to 5’UTR, leader 1, and leader 2 collectively as upstream sequences.

We decided to use the genotyped AIRR-seq data to characterize upstream sequence variants for all genes and alleles. To extract the upstream sequences, we removed the VDJ and constant regions, while keeping the original sequence’s V-region annotation. Sequences from each individual were processed separately. We observed slight variations in the length of 5’UTRs assigned to the same gene. It is important to have matching length for clustering, as different lengths could mean that identical sequences would not cluster together. To overcome this issue, for each gene we trimmed the ends of 5’ ends of the upstream sequences to match the most frequent length. We then took the trimmed upstream sequences with the same allele annotation and clustered them. Each cluster of a sufficient size gave rise to one consensus upstream sequence. This process was repeated for all genes and alleles across all individuals. Finally, consensus sequences from all individuals were combined to create an upstream germline reference dataset of the cohort (Fig.3). The number of individuals carrying each of the variants is shown in Supplementary Fig.5.

**Figure 3.**
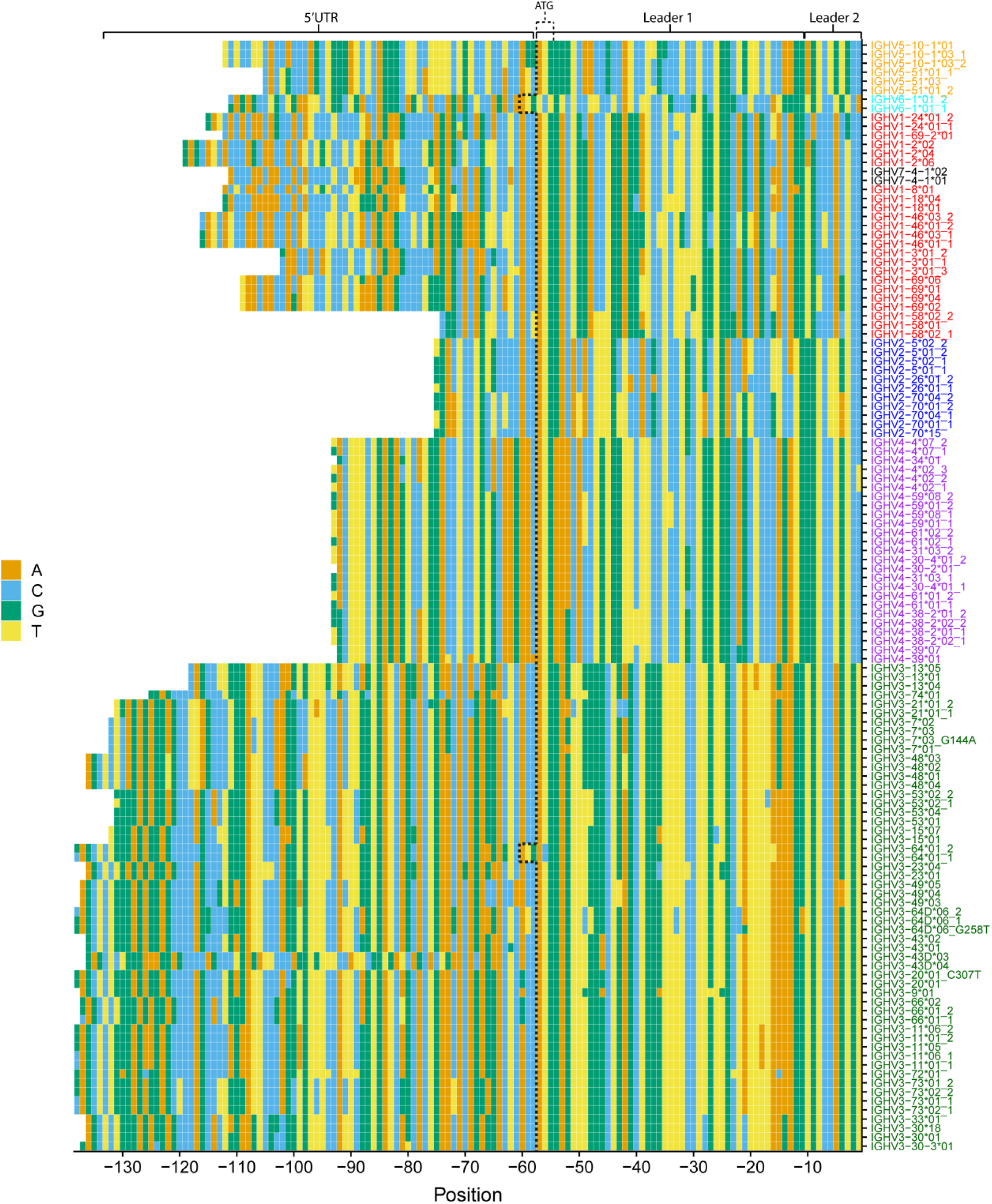
Upstream germline reference dataset. For each allele, consensus upstream sequences were built. Consensus sequences constructed from clusters with less than 10 sequences or with relative frequency < 0.1 were excluded. Each row represents a consensus upstream sequence of a V allele with 5’ to 3’ orientation. The colors of the tiles represent the different nucleotides. The coordinates on the x-axis describe the position of each nucleotide relative to the start of the V-region (5’ to 3’) and are therefore labeled as negative numbers. Alleles with more than one consensus sequence are marked with the allele name followed by an underscore and the respective consensus sequence number. For example, the two different consensus sequences for allele *IGHV3-64*01* are marked as *IGHV3-64*01_1* and *IGHV3-64*01_2*. The number of individuals who carry each variant are shown in Supplementary Fig.5.

According to the constructed germline reference dataset, the lengths of leader 1 (45 nt) and leader 2 (10 nt) sequences appear to be well conserved across most genes, with the exception of *IGHV3-64*01* and *IGHV6-1*01* (Fig.3). The leader 1 sequences of these two genes are 3 nt longer, which makes the position of ATG appear to be shifted upstream. The length of the 5’UTR is relatively conserved within the same gene family, however, there is a large variability across different families. Genes of the *IGHV2* family have the shortest 5’UTR, while the 5’UTRs of *IGHV3* genes are the longest.

Comparison of the consensus sequences in the cohort with the reference sequences obtained from the IMGT/GENE-DB^10^ revealed some discrepancies between our data and the reference database. For example, the IMGT reference sequence of the allele *IGHV5-51*01* has T at position −3 in leader 2, while the reference sequences of the other reference alleles have G at this position. However, in our data, all *IGHV5-51* alleles have G at position −3, as illustrated in Fig.3. Our observation of G at position −3 in *IGHV5-51*01* was validated by targeted amplification and Sanger-sequencing of *IGHV5-51*01* from a homozygous individual (Supplementary Fig.6).

### Length of 5’UTRs correlates with the distance between TATA-box and start codon

As depicted in Fig.3, the length of the 5’UTR differs between *IGHV* gene families, but is relatively conserved within a gene family. To investigate whether the different length of 5’UTRs among the different families had any correlation with the distance from the promoter elements, we decided to inspect the reference gDNA sequence from the IMGT database. We collected the available germline reference sequences of the upstream flanking regions of V-gene promoters from the IMGT/GENE-DB and aligned them to look for conserved patterns.

Using the sequences from the IMGT reference database, we determined the distance between the ATG start codon and the reference or putative TATA-box. We found that this distance varied greatly between different gene families. By comparing this distance to the 5’UTR length from the AIRR-seq data, we observed that the distance between the ATG and the TATA-box correlated with the length of the 5’UTR (Supplementary Fig.7). Sequences with longer ATG to TATA-box distance had longer 5’UTRs.

### Differences in the upstream sequences can aid allele annotation

The novel allele *IGHV3-64*05_G265C* was initially not validated by amplification of the gene *IGHV3-64*, as Sanger sequencing revealed only *IGHV3-64*02* in a selected individual carrying the suspected polymorphism, and with no sequence corresponding to allele *\*05* being present (Fig.4b). Originally, this allele was ambiguously annotated as deriving from either *IGHV3-64*05* or *IGHV3-64D*06*, as it differs by one nucleotide from each of these alleles (Fig.4a).

The upstream sequences of *IGHV3-64* and *IGHV3-64D* differ all across their length, including the 5’UTR, leader 1, and leader 2 (Fig.3). The upstream regions of the novel allele *IGHV3-64*05_G265C* are identical to those of *IGHV3-64D*, which indicated that this is indeed an allele of *IGHV3-64D* and not *IGHV3-64.* Therefore, we decided to amplify the gene *IGHV3-64D* using primers specific to the duplicated gene only. This resulted in the novel allele being finally validated (Fig.4c). Upon obtaining the full germline sequence of the novel allele, we observed that its intron matched the one of *IGHV3-64D* and not *IGHV3-64* (Fig.4d).

The genes *IGHV3-43* and *IGHV3-43D* are another example of duplicated genes with differences in the upstream sequences. Unlike the previous example, *IGHV3-43* and *IGHV3-43D* seem to have identical leader 1 and leader 2 sequences but differ in the 5’UTR (Fig.3). However, not only genes, but also some alleles of the same gene can be distinguished by their upstream sequences. The novel allele *IGHV3-64D*06_G258T* differs from *IGHV3-64D*06* in one position located in leader 1. Similarly, *IGHV4-39*01* and *IGHV4-39*07* have three differences within the 5’UTR; and the alleles *IGHV3-43*01* and *\*02* differ in one position within the 5’UTR.

## DISCUSSION

Our analysis of the naïve B cell immunoglobulin repertoire data from 98 individuals revealed several novel polymorphisms both in the coding and in the upstream sequences of *IGHV* genes. To our knowledge, we are the first to provide a comprehensive overview of upstream (5’UTR, leader 1, and leader 2) *IGHV* sequence variants in an AIRR-seq dataset. We managed to validate a number of novel alleles by targeted amplification of genomic DNA of the same individuals. In addition, we report the presence of G at position 318 instead of A in the gDNA sequence of *IGHV3-7*02*, which supports the findings of previous studies^31, 32^.

We faced several issues with missing or incomplete genomic reference sequences, which complicated the design of efficient primers for verification of novel alleles. Some of our validation attempts were unsuccessful resulting only in the amplification of a “wild-type” allele without a polymorphism. We suspect this might be caused by allelic dropout^33, 34^. As we show in our upstream sequence overview (Fig.4), alleles *IGHV4-39*01* and *IGHV4-39*\**07* differ at multiple positions within the 5’UTRs. Our primers were designed to bind flanking sequences of the gene, and their design was based on the current reference genome, which contains the allele *\*01* of *IGHV4-39.* Potential differences in the primer binding regions could be the cause of a failure to amplify the novel alleles, in this case *IGHV4-39*07_C288A*.

**Figure 4.**
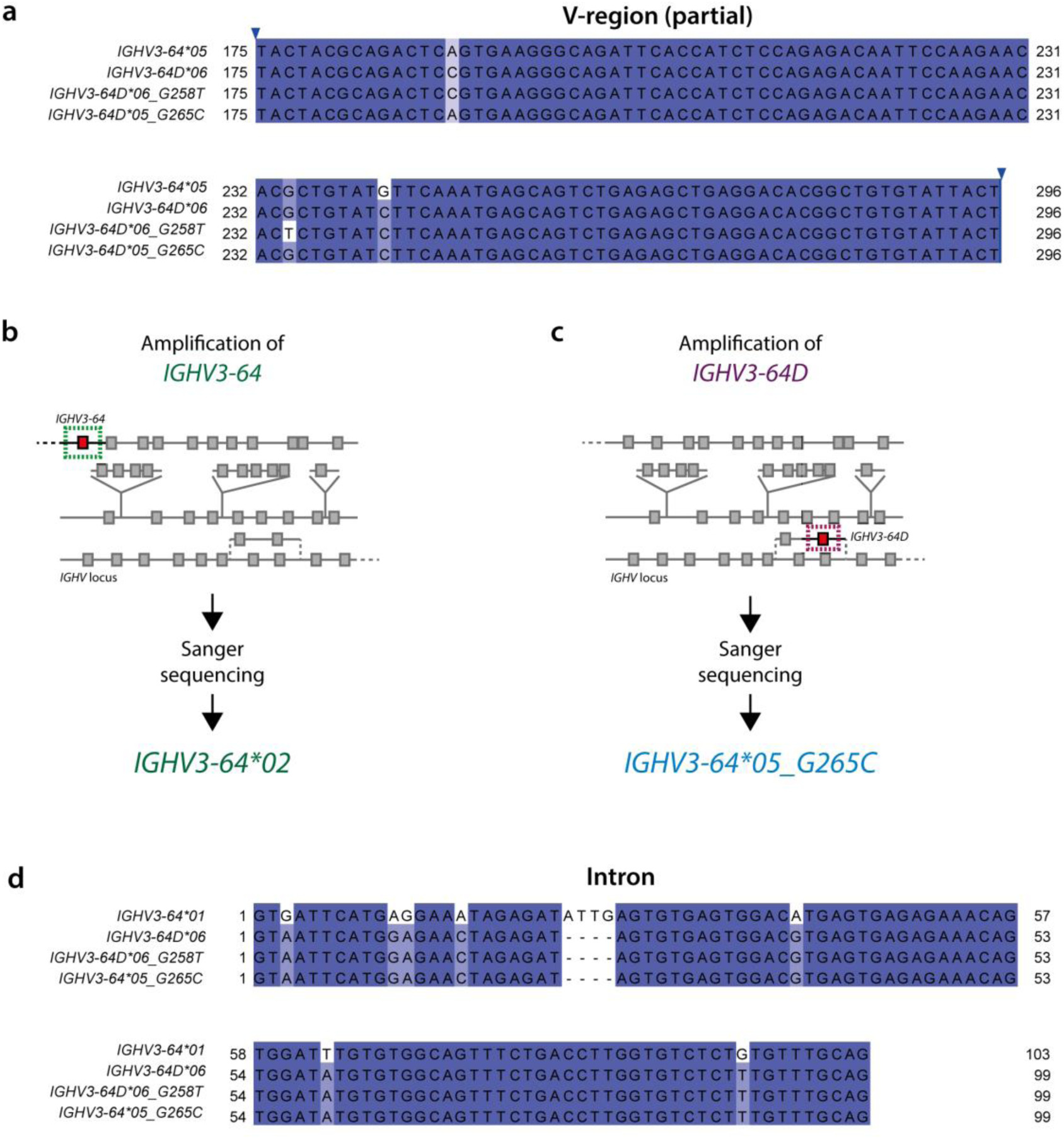
Genomic validation of *IGHV3-64(D)* alleles. (a) Alleles *IGHV3-64*05* and *IGHV3-64D*06* differ in only two positions within the V-region. To validate novel *IGHV3-64* and *IGHV3-64D* alleles found in the AIRR-seq data and ensure their correct annotation, we PCR amplified the genes *IGHV3-64* and *IGHV3-64D* from gDNA of selected individuals using gene-specific primers. (b, c) The process of validation of *IGHV3-64*05 G265C* depicted with a schematic *IGHV* locus representation. The novel allele was originally assigned as being closest to *IGHV3-64*05*, however, this allele was not amplified by primers specific for *IGHV3-64*. The novel allele was detected when *IGHV3-64D* was amplified. (d) Comparison of the intronic regions of *IGHV3-64*01*, *IGHV3-64D*06* and the novel *IGHV3-64D* alleles. The reference sequence of *IGHV3-64*05* in the IMGT database is partial and lacking the intron, and therefore could not be compared. The intron of the novel allele originally annotated as *IGHV3-64*05 G265C* matches the one of *IGHV3-64D*. The numbers in the alignments (a,d) do not follow the unique IMGT numbering.

Although AIRR-seq studies are very useful for characterizing variation in immunoglobulin genes, one of the main limitations are issues with gene and allele annotation^35^. The V-region is annotated based on the most similar allele in the reference database. However, since the V genes are highly similar, this annotation might not always be correct. Incorrect gene assignment could lead to potential downstream errors in analysis. In our study, the novel allele originally annotated as *IGHV3-64*05_G265C* was later found to be derived from the gene *IGHV3-64D*, located on a different part of the *IGHV* locus than *IGHV3-64*. As previously shown^4, 5, 9^, *IGHV3-64D* is likely a part of an alternative haplotype, since it was found to be deleted in many individuals, even in this cohort^30^. These two genes differ in their upstream sequences, and thanks to this distinction, we were able to correctly assign the novel allele to *IGHV3-64D* and validate it from gDNA.

Our results demonstrate that polymorphisms in the upstream regions can be utilized to improve annotation methods presently employed. Having said that, the genetic variation in the sequences upstream of the V-region is currently poorly characterized. Many reference sequences, which were deposited to the IMGT germline database are partial and contain only the V-region sequence. It is surprising that the genetic variation in the upstream regions is overlooked, considering the fact that the leader regions are frequently used as primer binding sites for immunoglobulin repertoire library preparation protocols^32, 36, 37^.

The reason for the existence of upstream polymorphisms is unclear, but conceivably such polymorphisms might have functional relevance by influencing stability of the mRNA or by affecting the binding of regulatory proteins^38, 39^. Further studies are needed to explore polymorphisms in the upstream sequences and to determine whether they have any functional effect. Association of these allelic variants with disease can be studied in sufficiently powered studies. In addition, more genomic studies could be performed to characterize their promoters and other regulatory elements, which might help explain the differences in expression levels across individuals.

## METHODS

### AIRR Sequencing of naïve B-cells

The data was obtained as a part of a previously published study^30^. In summary, naïve B cells from 100 individuals were sorted from peripheral blood mononuclear cells (PBMCs). The RNA was isolated and quality checked before being sent to AbVitro, Inc for library preparation and sequencing on Illumina MiSeq (2×300bp). About half of the cohort are celiac disease patients, and these subjects were included to increase the diversity of the cohort. Of note, this study was not designed and powered to perform comparative analysis of allelic frequencies between patients and controls.

### Amplification of target genomic regions

Genomic DNA (gDNA) was isolated from previously sorted T cells and monocytes (CD19-CD3+/CD14+)^30^ using the QiaAmp DNA mini kit (Qiagen), and the concentration was measured on Nanodrop.

Primers for validation were designed by PrimerBLAST using the reference genome as a template. The nucleotide sequences of primers with additional details can be found in the Supplementary material. For amplification of genes *IGHV3-7, IGHV3-20*, and *IGHV3-21*, primers from a recently-published study ^32^ were used. All oligos were synthesized and purified (RP-cartridge) by Eurogentec.

The target regions of the gDNA were amplified by touch-down PCR using Q5^®^ Hot Start High-Fidelity DNA Polymerase (NEB). Approx. 100 −200 ng gDNA from an individual with a suspected polymorphism was used as a template. The PCR started with two cycles with the annealing temperature of 70°C. The touch-down part of the PCR consisted of 10 cycles with the annealing temperature decreasing from 70°C to 60°C by 1°C every cycle. In the next 13 cycles, the annealing temperature remained constantly at 60°C, and the last step of the PCR was the final extension at 72°C. The length of the PCR product varied depending on the amplified gene, ranging between 750bp and 986bp.

### Cloning

The PCR products were cleaned using the Monarch^®^ DNA Gel Extraction Kit (NEB), and 3’ end A-overhangs were added by NEBNext^®^ dA-Tailing Module (NEB). The A-tailed products were subsequently cloned into pGEM^®^-T Easy vector (Promega) using the manufacturer’s protocol. For transformation, 4 µl of the ligation reaction were used to transform 90 µl XL10 CaCl2-competent cells. After transformation, 100 µl cells were plated on LB_amp_ 50 µg/ml plates that have been previously coated with IPTG/X-Gal (40 µl 100 mM IPTG + 16µl 50 mg/ml X-Gal). The IPTG/X-Gal treatment allows for selection of successfully transformed colonies based on color. After overnight incubation at 37°C, white colonies were picked and the plasmids were isolated using the Monarch^®^ Plasmid Miniprep Kit (NEB). To verify that the picked colonies contain an insert of the correct size, a PCR was performed using the same primers as for the amplification of gDNA, and the products were analyzed by gel electrophoresis (1% agarose, 100 V, 35 min). The size of the PCR product was between 750-986bp, depending on the gene amplified.

### Sanger sequencing

Sanger sequencing of the plasmid DNA containing the correct-sized insert was performed by Eurofins. The resulting sequences were trimmed to remove the vector and primer sequences. V-gene annotation was done by IMGT/HighV-QUEST ^40^. To check for polymorphisms in the introns, leader regions and 5’UTRs, the trimmed sequences were aligned by MUSCLE ^41, 42^ to the reference alleles of the amplified gene, where available, and checked for polymorphisms. Alignments were visually inspected in Jalview^43^ and/or UGENE^44^.

The sequences were named based on the amplified gene, followed by the closest reference allele and the V-region polymorphism, which was determined by IMGT V-Quest ^45^ or by manual annotation (in cases of ambiguous annotation).

The gDNA sequences of validated novel alleles were submitted to GenBank and subsequently to IMGT.

### AIRR-seq data pre-processing

The AIRR-seq data was pre-processed as described originally^30^ using pRESTO ^46^. Two individuals were excluded from the analysis due to low sequencing depth (<2000).

### Novel allele discovery and genotype inference

Genotype inference and novel allele discovery was also performed by TIgGER v 0.3.1 and IgDiscover v0.11. The pre-processed sequences were annotated by IgBLAST 1.14.0^47^ with modified parameters, and the IMGT germline database (24) from January 2019 was used as a reference. The results of alignment and genotype inference by TIgGER were processed using a similar pipeline to the one used in http://www.vdjbase.org with slight modifications.

We experienced that the default settings resulted in incorrect annotation for some genes. This was particularly obvious for the allele *IGHV5-51*03*, which was incorrectly annotated as *IGHV5-51*01* with one mutation C45G, corresponding to the already known allele *\*03*. These two alleles differ only by one nucleotide, and it was the length of the reference allele that seemed to affect whether or not the sequence was correctly annotated by IgBLAST. The reference for *\*03* is 2 nt shorter than the reference sequence for *\*01*, while sequences in our data corresponding to *IGHV5-51*03* were matching the length of allele *\*01*. Adjusting the IgBLAST parameters --reward to 0 and --penalty to −3 resolved this annotation problem. These parameters were also induced manually in IgDiscover alignment step.

For novel allele detection we tested the parameters of the TIgGER function “findNovelAlleles”: 1) germline_min to 50,100 and 200 (default). 2) j_max to 0.15 (default), 0.3 and 0.5. 3) min_seqs to 25 and 50 (default). Different parameters resulted in different sets of novel alleles identified. To allow for discovery of novel alleles in lowly-expressed genes, we set the germline_min parameter to 50. The rest of the parameters, including j_max and min_seqs, was left as default. The novel alleles were further submitted for genotype inference, using a Bayesian approach, for each individual. As for IgDiscover, the default parameters for novel allele and genotype calls were applied. Analysis of the IgDiscover and TIgGER output was performed in R Studio version 3.6.0.

### Filtering out false positive suspects

Errors that occur during the PCR reaction and/or sequencing could result in a false novel allele call. To filter out the suspected false positive signals, we first determined the mismatch frequency for all novel allele candidates. Novel allele candidates with low mismatch frequency were considered as false positives. These included all alleles with mutation patterns A152G, T154G, and A85C. Although the mismatch frequencies of sequences with the A85C polymorphism seemed to follow a bimodal behavior (Supplementary Fig.1), the higher frequency mode that should correspond to heterozygous individuals is centered around 20%, and not 50% as would be expected. As a result, they were not considered as true novel alleles. On top of that, this polymorphism was only observed in four individuals that were sequenced in a pilot separately from the other samples, and A to C mutation is the most common substitution error in Illumina MiSeq^48^.

### Analysis of gene and allele usage

Following the inference of genotype for each individual, we used IgBLAST 1.14.0^47^ to re-align each individual’s sequences with their own personalized germline *IGHV* database as inferred by TIgGER. To compare the relative gene usage in individuals with different allele combination, we selected sequences with V-region length >200 and up to 3 mutations. Since the duplicated genes *IGHV3-23*01* and *IGHV3-23D*01*; *IGHV1-69*01* and *IGHV1-69D*01*; *IGHV2-70*04* and *IGHV2-70D*04* have identical V-regions, they often result in ambiguous allele assignment. Annotation for sequences with ambiguous allele assignments for these genes were renamed *IGHV3-23*01D*, *IGHV1-69*01D* and *IGHV2-70*04D*, respectively. Additionally, *IGHV3-30-5*01* and *IGHV3-30*18* are also identical; and we renamed them as *IGHV3-30X*doub*; and *IGHV3-30X*trip* if the sequence annotation also contained *IGHV3-30*01* as another possible assignment. All remaining sequences with multiple allele annotations were filtered out. To plot the relative gene usage, we first calculated the relative usage fraction of each allele of a gene separately. Afterwards, we summed up the relative usage fractions of alleles of the same gene and plotted the relative usage of each gene across all individuals.

### Inference of upstream sequences (5’UTR, leader 1 and leader 2)

We decided to look at the upstream regions that consist of (5’-3’) 5’UTR, leader 1, and leader 2. For the analysis of the upstream regions, only sequences with up to 3 mutations in the V-region (after novel allele inference and genotyping) and single assignment V-call were selected. For each individual, the V-region sequences were trimmed away and the remaining upstream sequences of the same V-gene were aligned by the last nucleotide of leader 2 sequence and flipped 3’-5’.

Since the length of the 5’UTR sequences of the same gene in AIRR-seq data can vary due to whole VDJ sequence length and sequencing length limitations, we needed to determine where to trim the longer sequences. To do this, we first filled the ends of sequences with Ns to match the length of the longest sequence for the respective gene. We then trimmed all sequences after the first position, at which 95% sequences contained N.

After that, for each allele and for each individual, we removed all artificially added Ns. Next, we estimated sequence lengths, and lengths with frequency above 0.1 were considered frequent. Sequences shorter than the shortest frequent sequence length were filtered out and sequences longer than the longest frequent sequence length were trimmed to match its length. By applying ClusterSets.py (--ident 0.999, --length 0.5) and BuildConsensus.py (--freq 0.6) from pRESTO, we constructed clusters that resulted in consensus sequences for each allele. For each cluster we calculated its frequency based on the number of sequences assigned to it. Clusters with frequency below 0.1 or with less than 10 sequences were removed.

For each allele, consensus sequences from all individuals, were trimmed to match the shortest consensus sequence, and identical sequences were re-collapsed by allele and individual. For some of the consensus sequences, one of the nucleotides was marked with ambiguous assignment (N) by BuildConsensus.py function. In such cases, the original cluster was split into two clusters based on the ambiguous assignment and consensus sequences were reconstructed manually. Finally, to create the consensus upstream sequences, for each allele the trimmed sequences were submitted to ClusterSets.py (--ident 1.0, --length 1.0) and BuildConsensus.py (--freq 0.6) functions and as a result, for each gene and allele a set of consensus V upstream sequences were gathered. In the last step, we compared and collapsed identical sequences from all individuals to create a database of upstream sequences in the cohort.

### Analysis of the reference germline upstream sequences

Reference germline sequences of the upstream sequences, including the 5’UTR, were obtained from the IMGT GENE-DB and by searching through the IMGT “Gene tables” in order to get an alternative longer sequence if available. The reference upstream sequences longer than 150 nt were aligned using the MUSCLE tool at EMBL-EBI ^42^, and the alignment was visualized by Jalview ^43^ to look for conserved regions. The obtained consensus sequences of conserved regions were compared to IMGT resources for annotation. The TATA-boxes were determined based on either the reference annotation by IMGT, searching through previous studies, or by looking for a TA-rich region downstream of the octamer. Promoters studied by older studies include that of *IGHV6*^49^ (with two TATA-boxes) and *IGHV1*^50^.

*IGHV2* analysis is based on the available upstream reference sequences of *IGHV2-5*01,*02* and *IGHV2-70D*04,*14*. *IGHV3* schematic promoter representation was based on the upstream reference genomic sequences of *IGHV3-43*01, IGHV3-48*02, IGHV3-49*03, IGHV3-64*02, IGHV3-64D*06* and the genomic sequences obtained by Sanger sequencing of *IGHV3-7*02* and *IGHV3-64D*06*. The IGHV4 schematic representation of the promoter was based the reference genomic sequences of *IGHV4-4*07* and *\*08*; *IGHV4-28*01,*02,*07*; *IGHV4-30-2*06; IGHV4-30-4*07; IGHV4-31*02; IGHV4-34*01,*02,*11; IGHV4-38-2*02; IGHV4-39*01; IGHV4-59*01,*02,*11;* and *IGHV4-61*01,*08,*09*.

### Data availability

The pipeline for novel allele discovery and genotype processing using the software tools TIgGER and IgBLAST is available on the VDJbase website (https://www.vdjbase.org). Custom code for the analysis of upstream sequences is available at https://bitbucket.org/yaarilab/cluster_5utr/src/master/.

Sanger sequences of validated *IGHV* alleles have been deposited in the GenBank under accession numbers: MN337615 (*IGHV1-2*02_G207T*), MN337616 (*IGHV1-3*01_T35A*), MN337617 (*IGHV1-46*01_C213G*), MN337618 (*IGHV1-69*01_C243T*), MN337619 (*IGHV3-7_G144A_T300C*), MN337620 (*IGHV3-7*02_A318G*), MN337621 (*IGHV3-9*01_T307C*), MN337622 (*IGHV3-20*01_C307T*), MN337623 (*IGHV3-21*01_C255T*), MN337624 (*IGHV3-64D*06_G258T*), and MN337625 (*IGHV3-64D*06_C210A*).

## ACKNOWLEDGEMENTS

This work was supported by grants from the Research Council of Norway through its Centre of Excellence funding scheme [project number 179573/V40], the South-Eastern Norway Regional Health Authority [project 2016113] and Stiftelsen KG Jebsen [SKGMED-017] to L.M.S., and grants from ISF [grant number 832/16] to G.Y., M.G. and A.P.

This project has also received funding from the European Union’s Horizon 2020 research and innovation program under grant agreement No 825821. The contents of this document are the sole responsibility of the iReceptor Plus Consortium and can under no circumstances be regarded as reflecting the position of the European Union.

Elements of Figure 1 were modified from Servier Medical Art, licensed under a Creative Common Attribution 3.0 Generic License (http://smart.servier.com/).

We would like to thank Knut E. A. Lundin for coordinating collection of blood samples of participating subjects and for being responsible for the ethical approval for the project. We also thank Victor Greiff for discussions and helpful advice; and Marie K. Johannesen and Bjørg Simonsen for technical assistance. We would also like to express our gratitude to all study participants.

## AUTHOR CONTRIBUTIONS

L.M.S. and G.Y. conceived and supervised the project; I.M., I.L. and O.S. carried out the experimental work; M.G., I.M., A.P., and G.Y. analyzed the data; I.M., M.G., and L.M.S. wrote the paper. All authors edited the manuscript.

## COMPETING INTERESTS

No conflict of interests declared.

## Supplementary Material

**Supplementary Figure 1.**
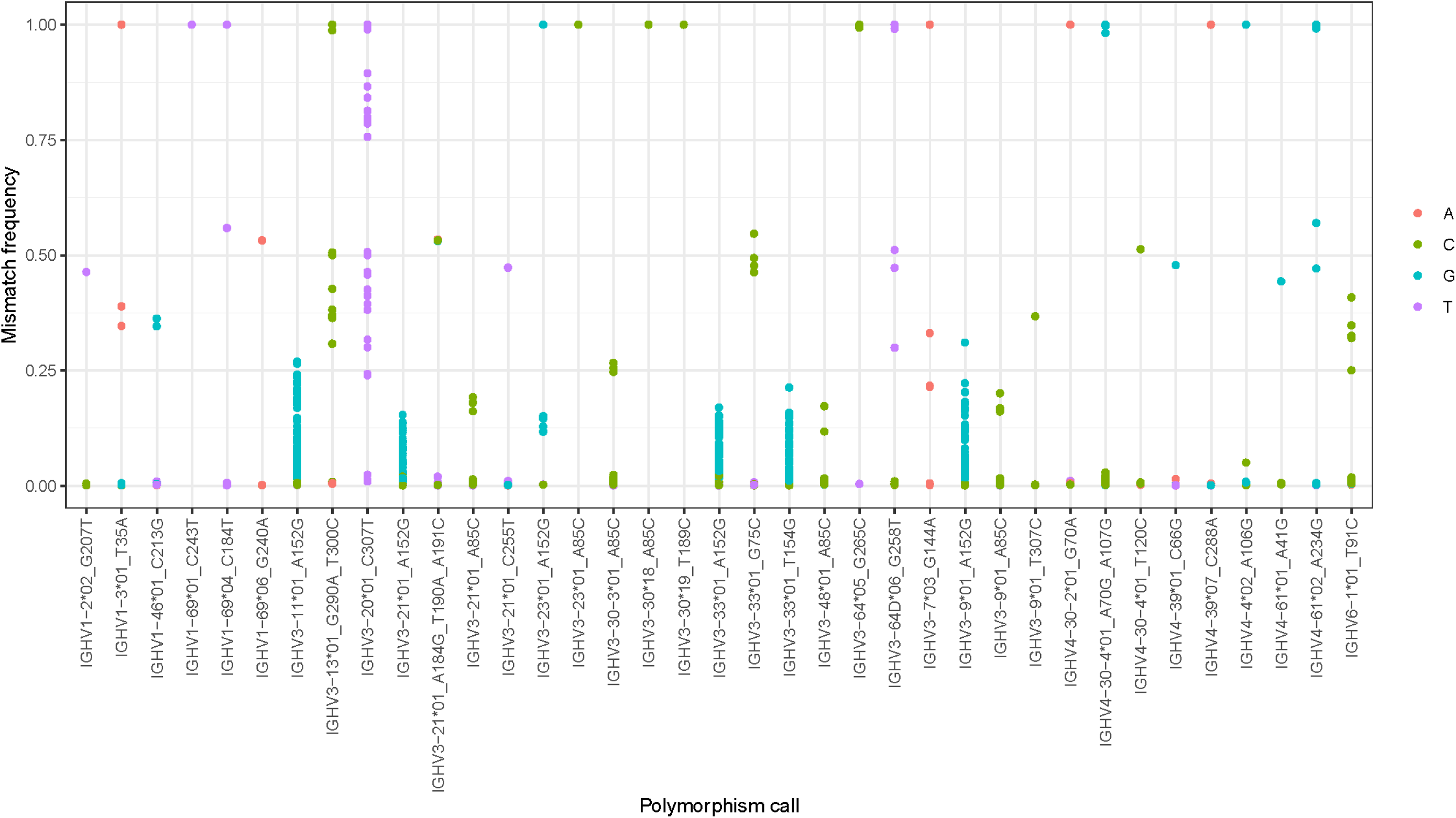
Germline mismatch frequency. For each individual, relative frequencies of polymorphisms (y-axis) were calculated for positions in sequences aligned to an allele, for which a novel allele was inferred in the dataset (x-axis). Each dot represents a mismatch frequency for an individual for a certain allele and nucleotide. The color of the dot represents the nucleotide that does not match the germline.

**Supplementary Figure 2.**
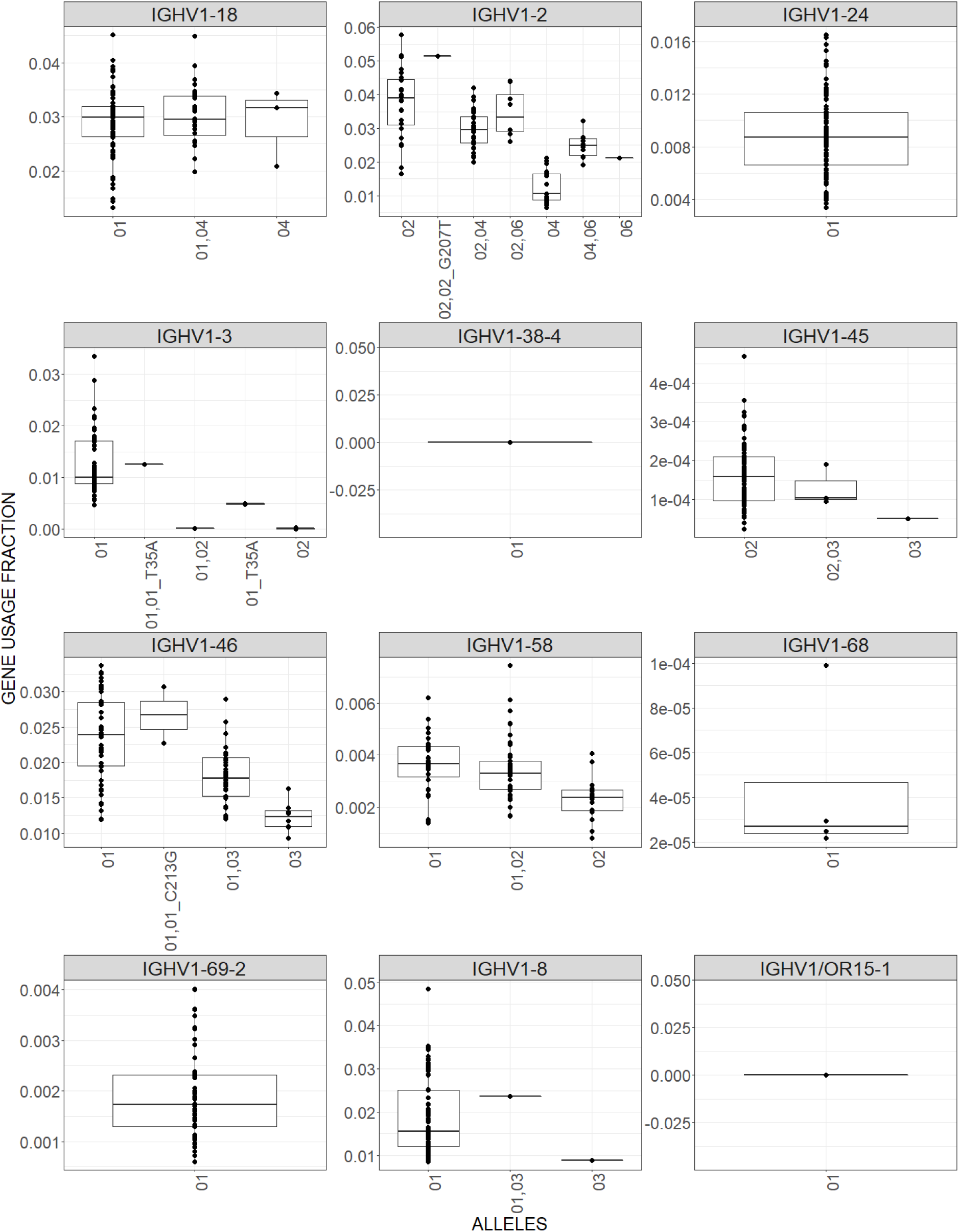
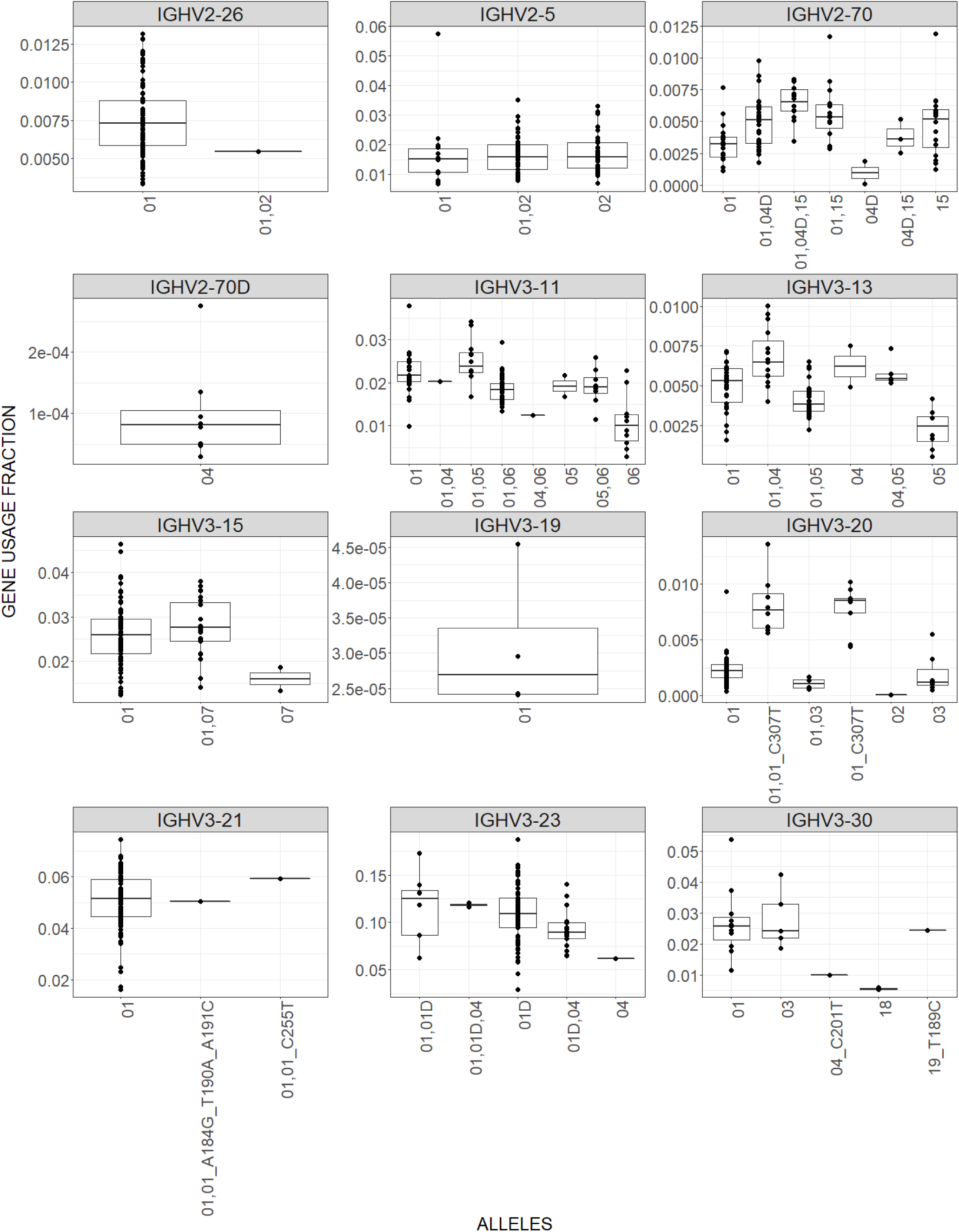
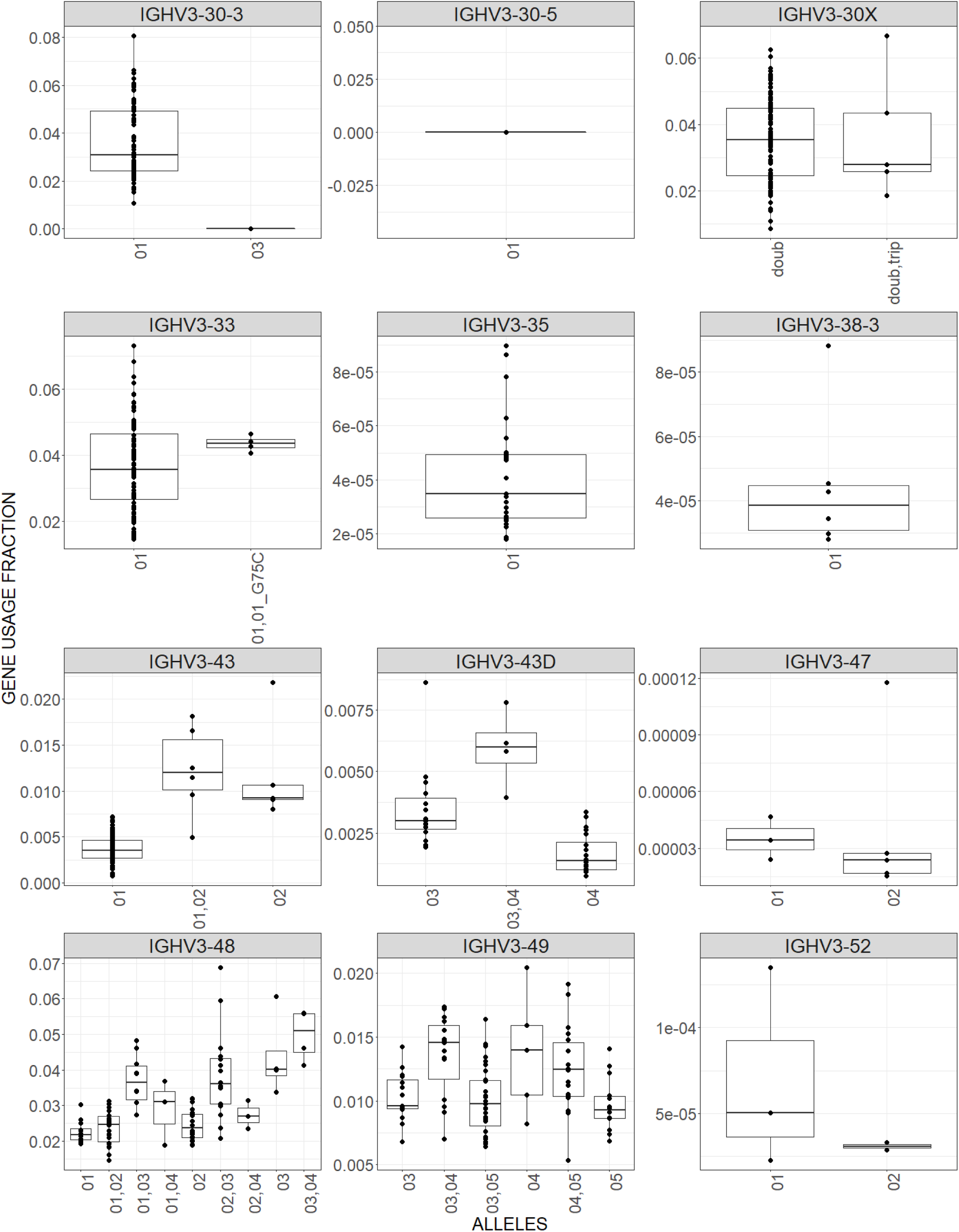
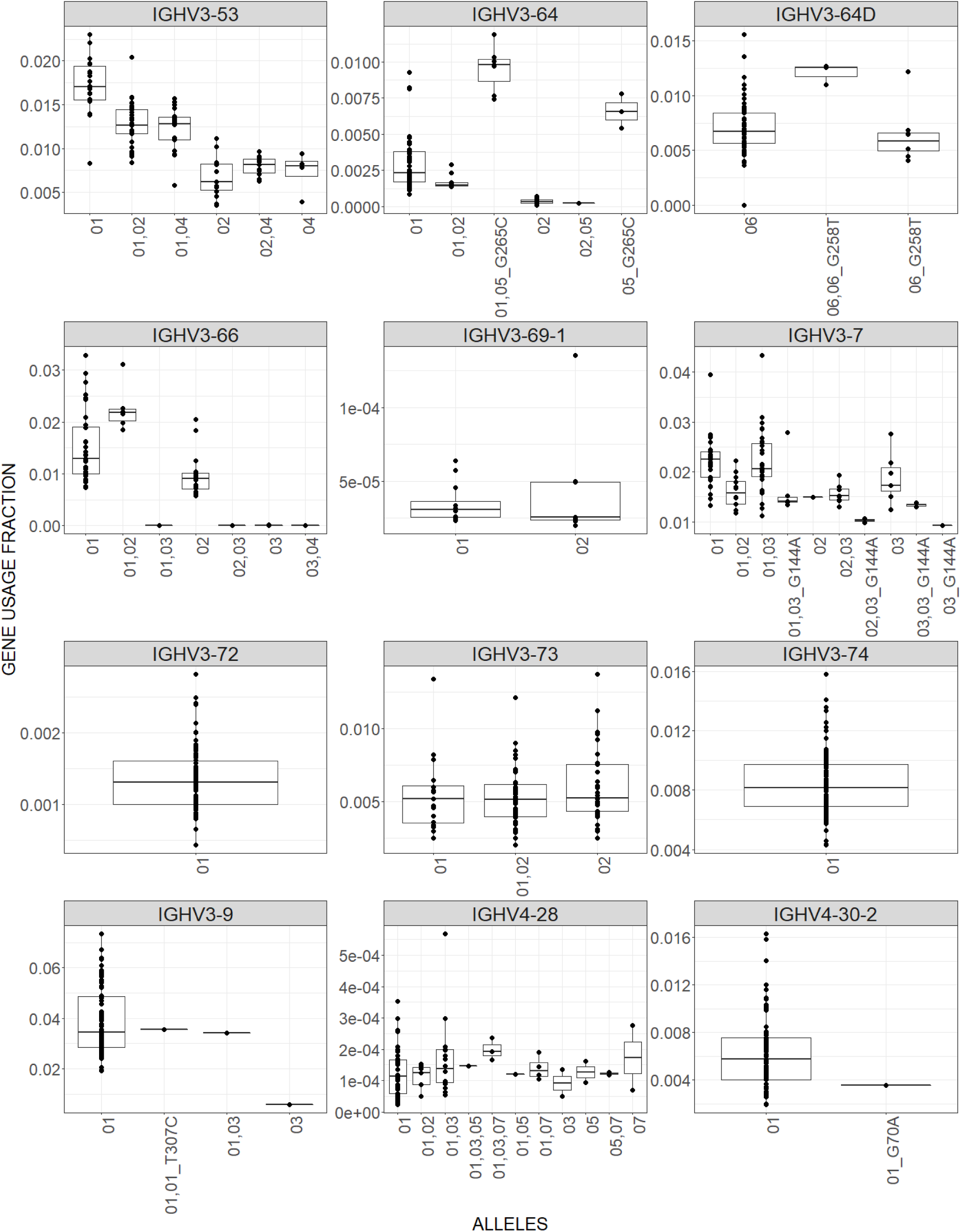
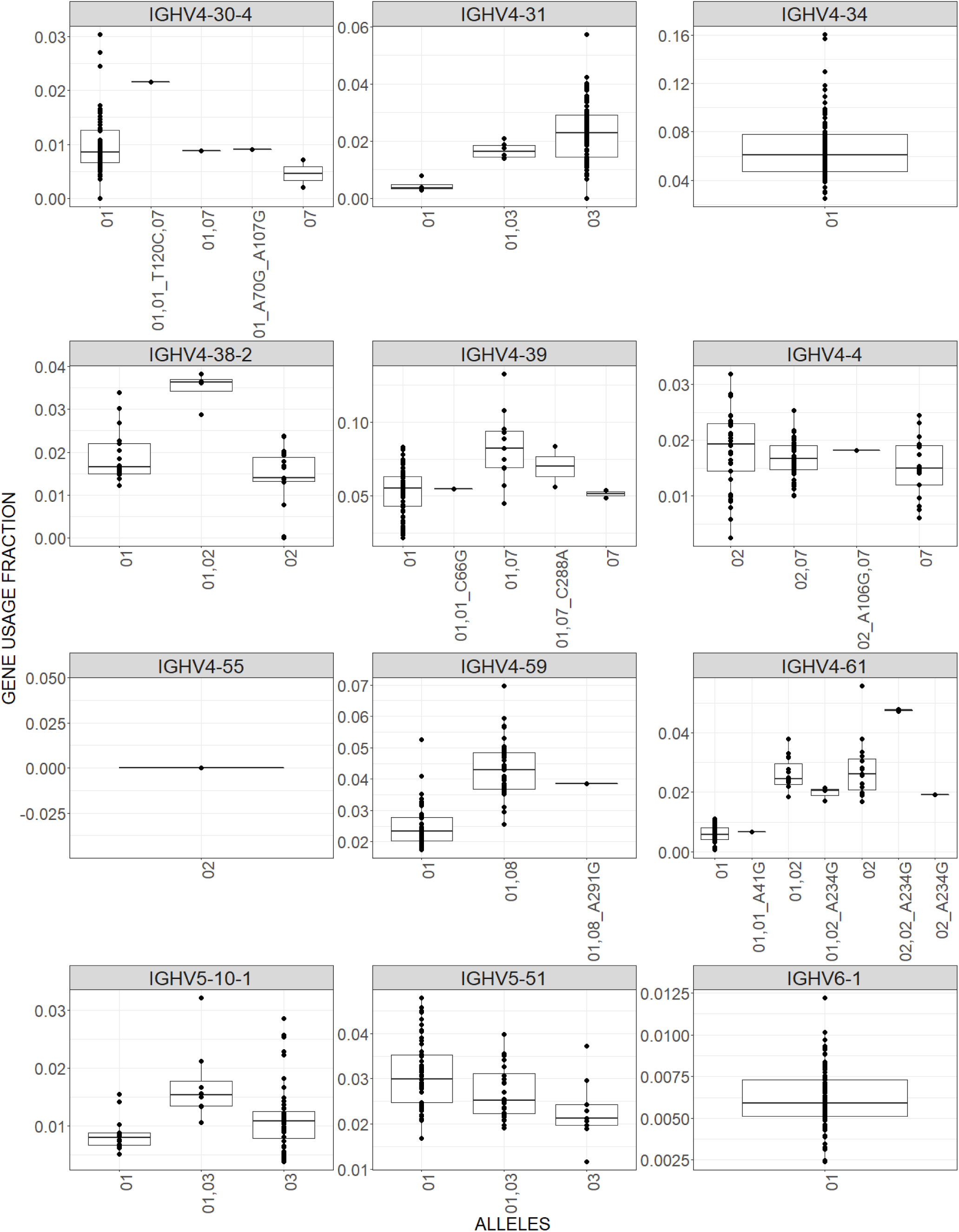
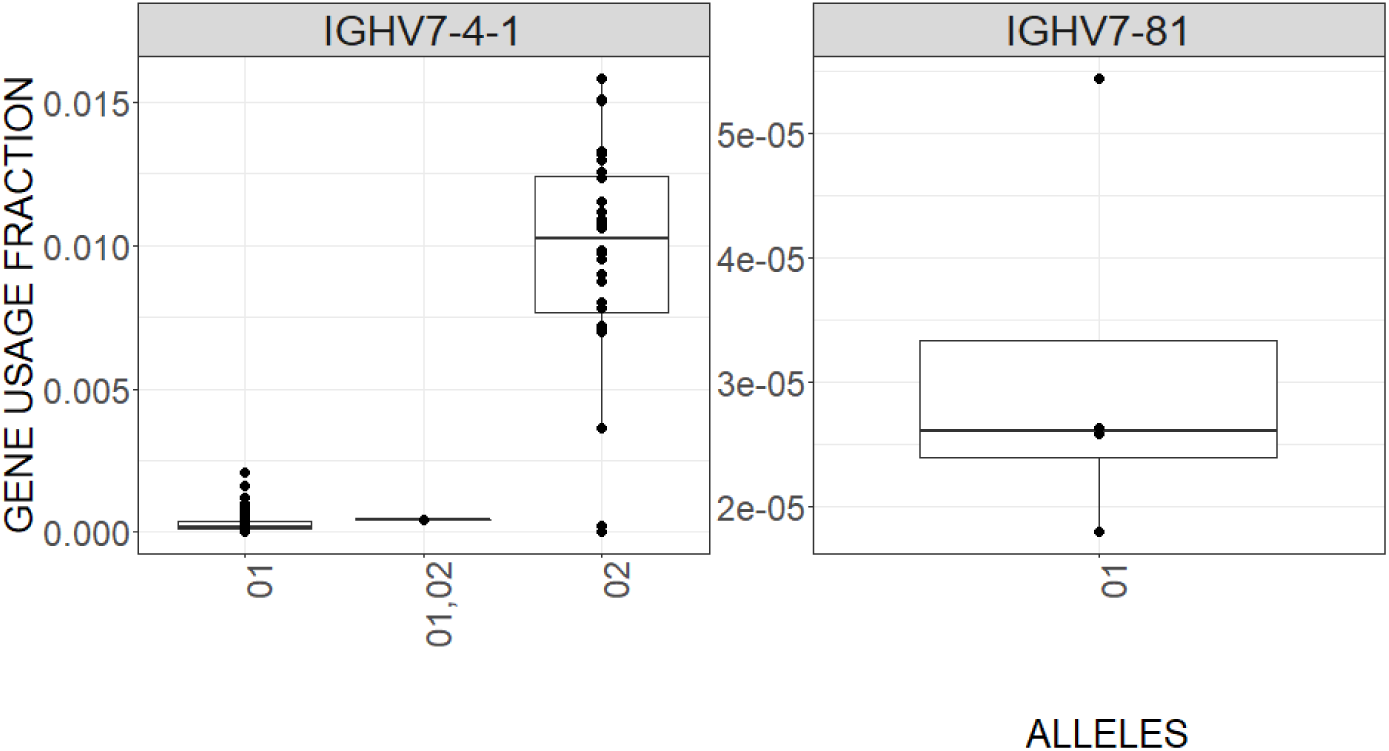
Usage of genes across individuals in the cohort. For each allele of a gene, we calculated its relative usage fraction in each individual. The usage fractions of alleles of the same gene in the same individual were then summed, revealing the gene’s usage fraction. The x-axis shows the inferred allele, or multiple alleles, that were found in an individual’s inferred genotype. Each dot represents one individual. The y-axis shows the relative usage fraction of a gene within the expressed repertoire. The bar represents the median value. A bias can be observed in some genes, where the median gene usage is higher in individuals homozygous for a specific allele than those homozygous for another allele. This figure continues on the next five pages.

**Supplementary Figure 3.**
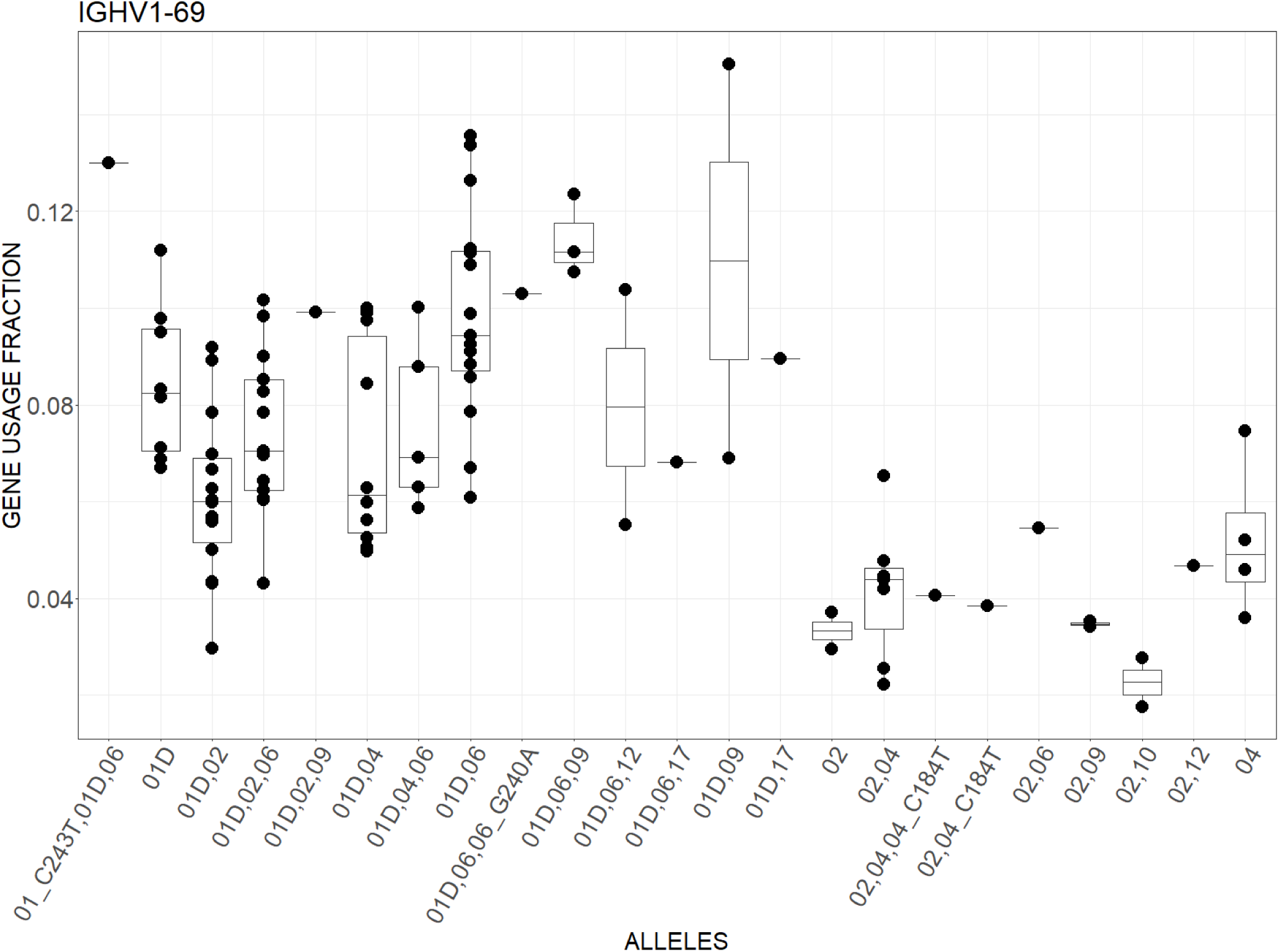
Usage of *IGHV1-69* across individuals in the cohort. Relative usage fraction was calculated for each allele separately and in each individual, and the relative fractions of all expressed alleles were summed up. Different combinations of expressed alleles are shown on the x-axis, and the summed gene usage fraction is shown on the y-axis. Each dot represents one individual. The bar in the boxplot represents the median value.

**Supplementary Figure 4.**
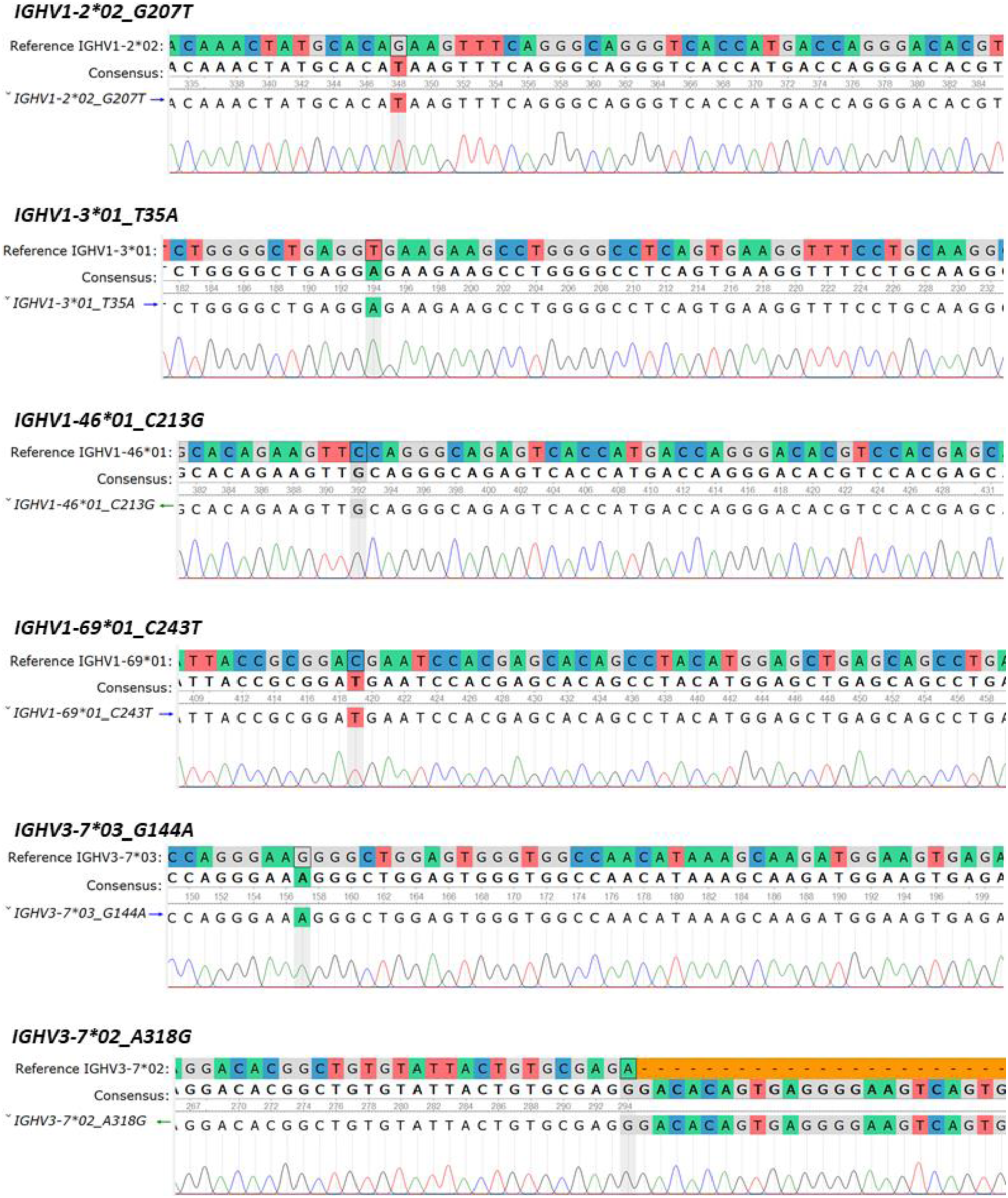
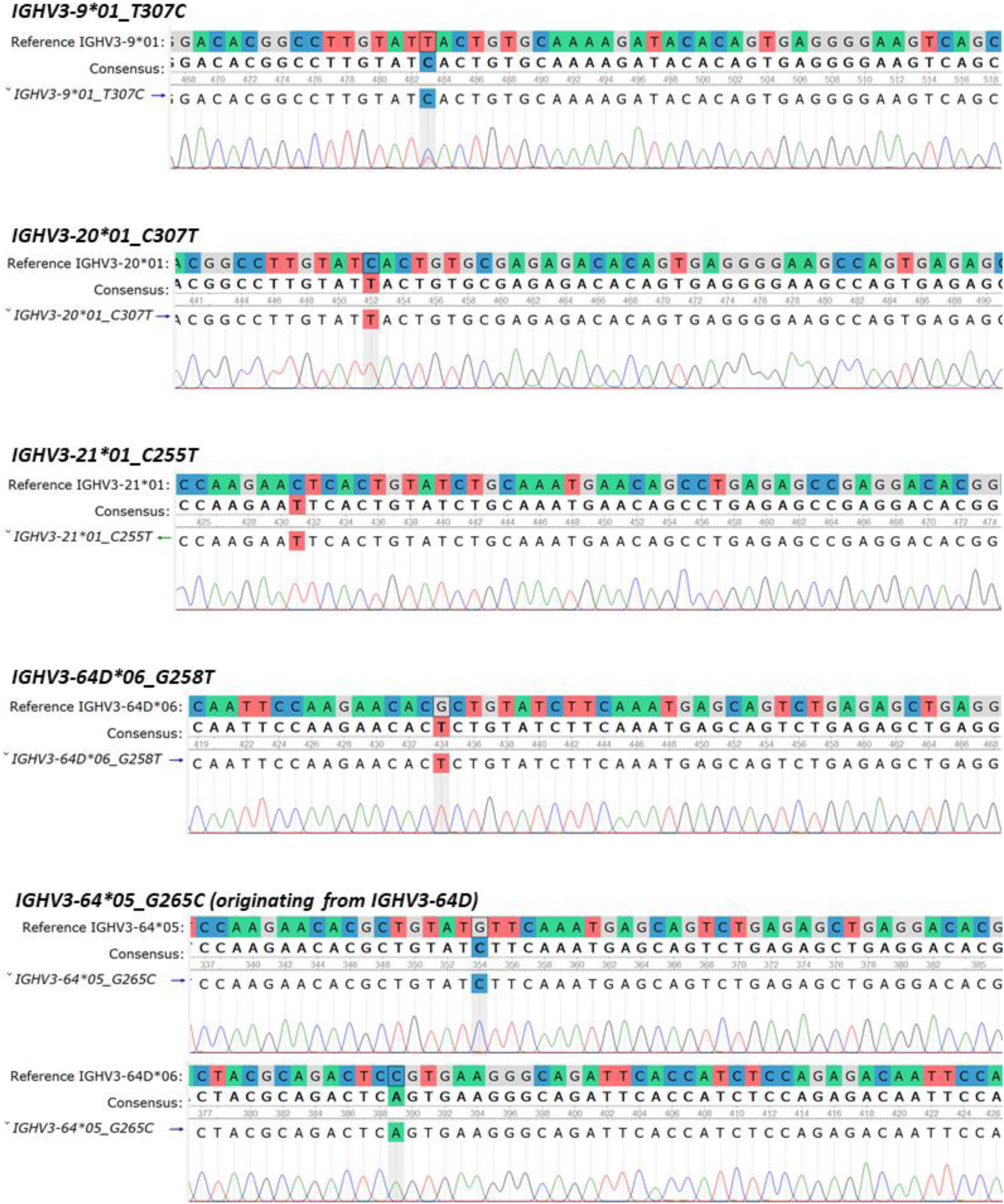
Sanger sequencing results. Ten novel alleles were validated by targeted amplification and subsequent Sanger sequencing. The trace files were aligned to reference sequences from IMGT GENE-DB^1^ and visualised by UGENE^2^. This figure continues on the next page.

**Supplementary Table 1.**
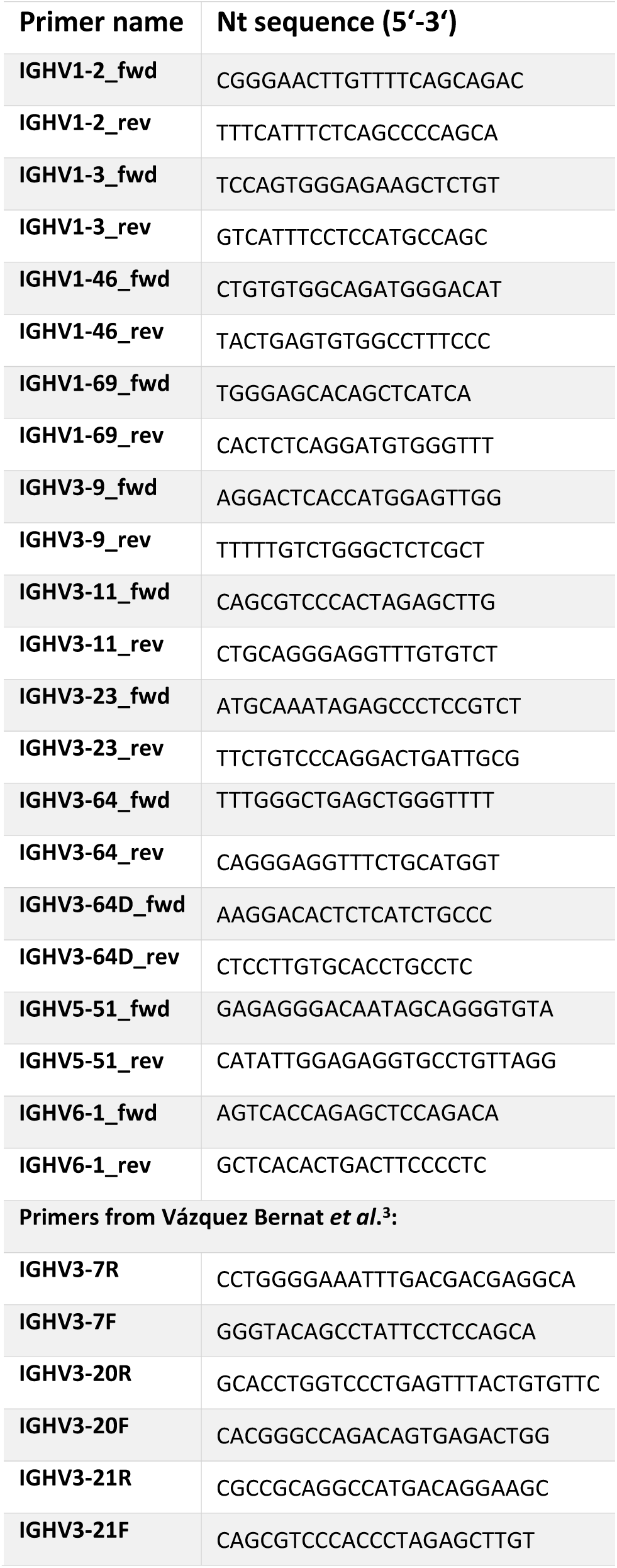
Primers used for genomic validation.

**Supplementary Figure 5.**
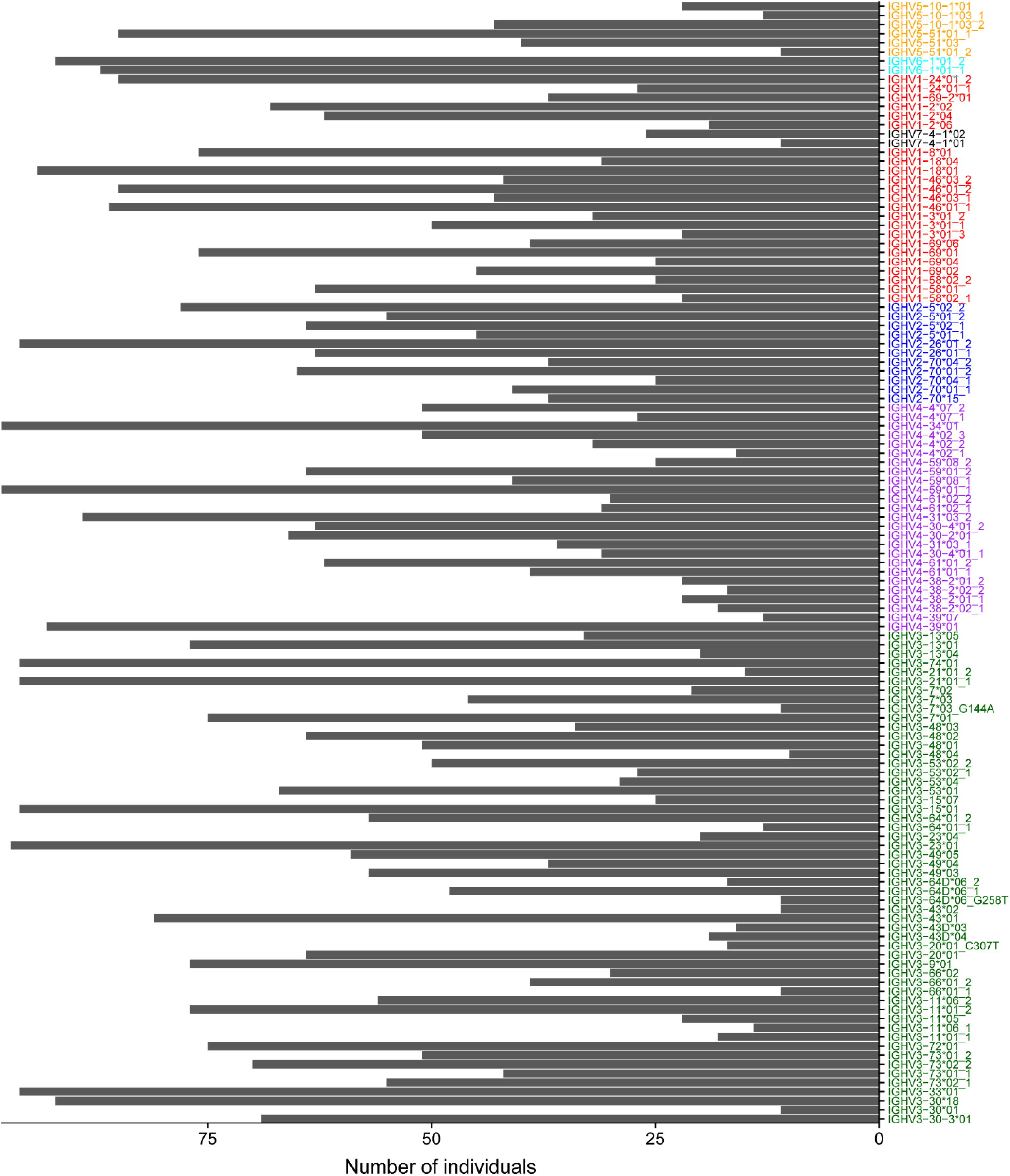
Number of individuals carrying the consensus 5’UTR sequence. Consensus 5’UTR sequences for each *IGHV* allele across all individuals were gathered and clustered to create a 5’UTR database. The length of each bar (x-axis) is the cluster size for a specific 5’UTR allele (y-axis), i.e. the number of individuals carrying the variant.

**Supplementary Figure 6.**
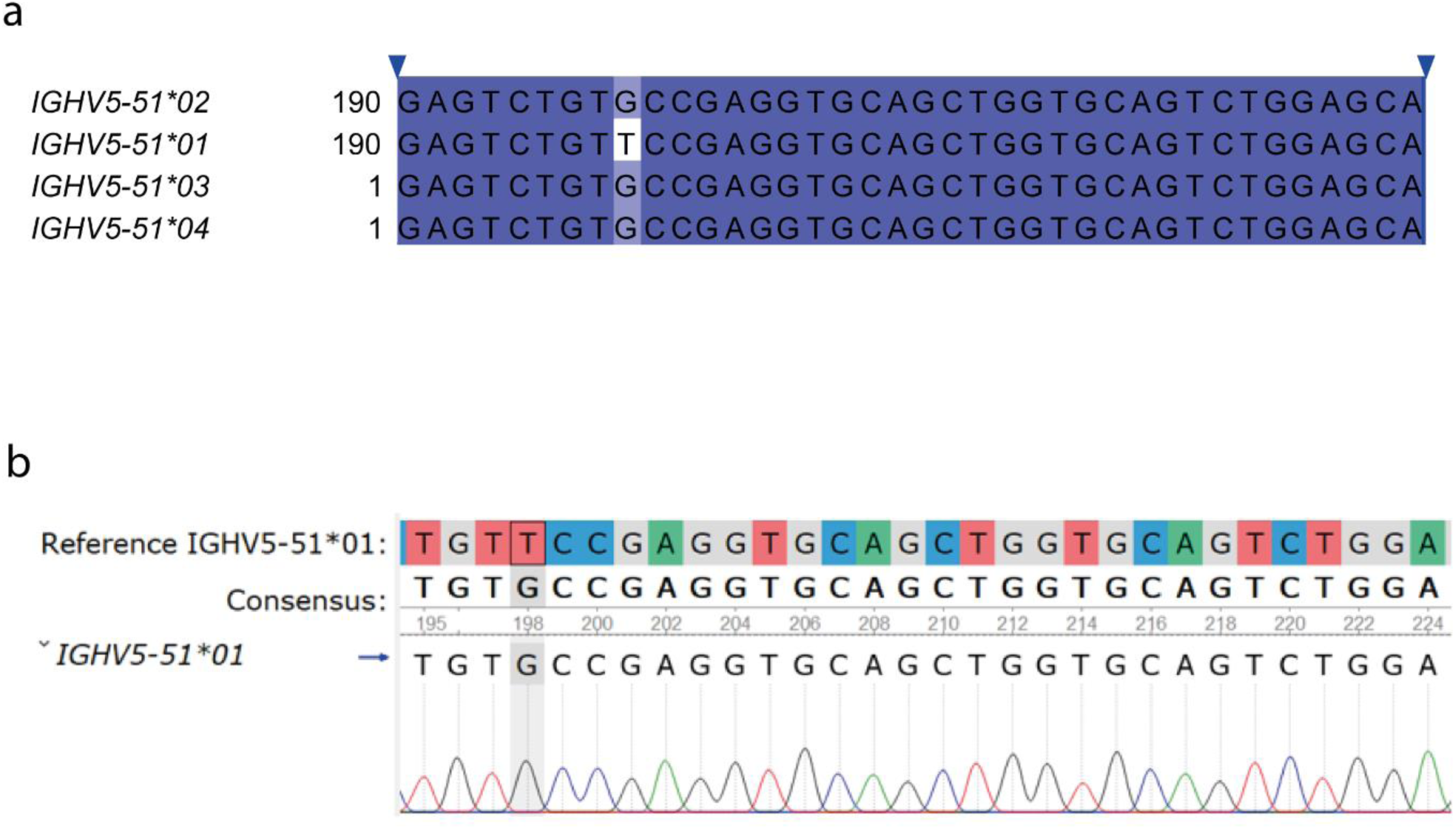
Validation of the *IGHV5-51*01* leader 2. The leader 2 part of *IGHV5-51*01* in all individuals in our cohort differed from the reference in the IMGT database. (a) Alignment of the reference leader 2 sequences of selected *IGHV5-51* alleles. (b) An individual from our cohort homozygous for *IGHV5-51*01* was selected and *IGHV5-51* was amplified using gene-specific primers (shown in Supplementary Table 1). Sanger sequencing of the amplified product revealed that the leader 2 of *IGHV5-51*01* indeed differs from the reference.

**Supplementary Figure 7.**
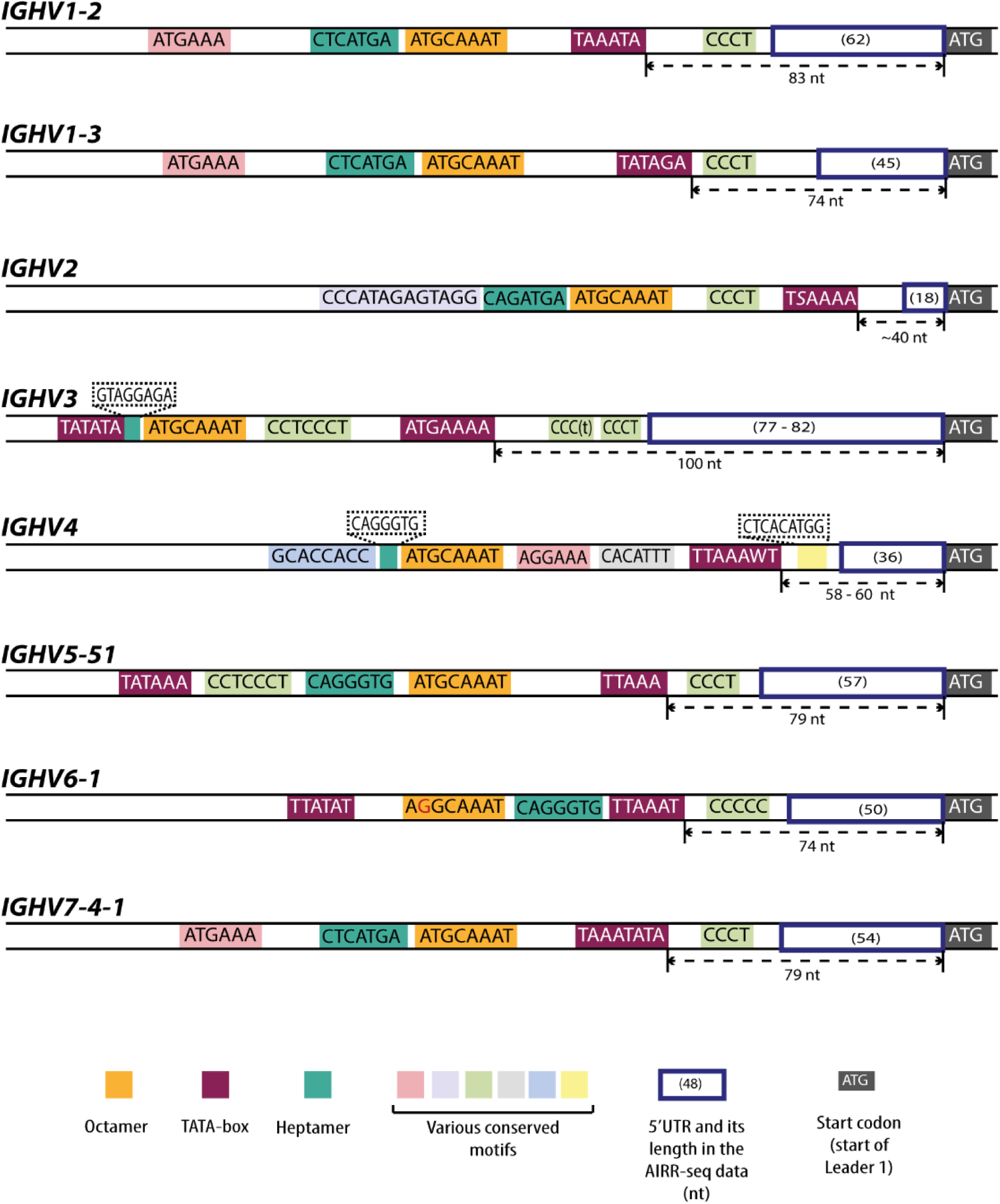
Schematic representation of the *IGHV* promoter regions. Reference upstream genomic sequences, including the promoter region were retrieved from the IMGT germline database and schematically depicted. Conserved motifs were identified by aligning all available 5’UTR and promoter reference sequences (> 150 nt) by MUSCLE and by searching for regions with high levels of homology. TATA-box sequences (in maroon) of some genes have been previously reported. For the remaining genes, we identified a putative TATA-box by searching for a TA-rich sequence. The octamer (in yellow) is well characterized and highly conserved across all genes. The heptamer (in dark turquoise) was only characterized for *IGHV1* genes. In the other genes, we identified putative heptamers by searching for a conserved sequence upstream of the octamer. Various conserved motifs with unknown function were also identified (pastel colors). The ATG start codon is shown in grey. The 5’UTRs that are found in the AIRR-seq data are lined in dark blue, and their typical length in the repertoire sequencing data is shown in brackets. The length of the 5’UTRs correlated with the distance between the ATG and the TATA-box.

